# Modeling CAR Response at the Single-Cell Level Using Conditional Optimal Transport

**DOI:** 10.1101/2024.11.11.622906

**Authors:** Alice Driessen, Jannis Born, Rocio Castellanos Rueda, Sai T. Reddy, Marianna Rapsomaniki

## Abstract

Chimeric Antigen Receptor (CAR) T cell therapy is a promising area of cancer immunotherapy. However, many challenges such as loss of persistence, T cell exhaustion, and therapy associated toxicities hamper further advancement of CAR T cell therapy. Therefore, recent efforts have focused on designing improved CARs that show better therapeutic characteristics. However, it is unfeasible to test all CAR variants in lab-based assays as CARs consist of multiple intracellular signalling domains. This results in over 100’000 possible variants. We leverage computational modeling to navigate this vast combinatorial space by learning the relationship between CAR design and T cell functionality, thereby proposing promising CAR T cell designs. CAR T cells expressing different variants can be viewed as cells that underwent different perturbations. Neural Optimal Transport is an upcoming field that can model single cell perturbations and predict unseen cells and conditions. In this work we leverage the conditional Monge Gap to model the response to CAR expression at a single-cell level and generate gene expression of cells that express an unseen CAR design. We show that CAR OT (CAROT) significantly outperforms the baseline for gene expression prediction for in-distribution CAR variants, with distinct gene expression patterns per CAR that capture biological characteristics. When predicting unseen CAR variants, we demonstrate promising results in terms of gene expression prediction and show the model learns gene expression patterns linked to domains in the training set. This work demonstrates that optimal transport may support discovery and development of new CAR T cell designs.

## 1 Introduction

Chimeric Antigen Receptor (CAR) T cell therapy is a promising area of cancer immunotherapy, with currently six FDA-approved therapies and over a thousand ongoing clinical trials [1]. CARs are synthetic protein receptors consisting of an extracelluar domain linked to an intracellular signalling domains, which carry the necessary T cell activation signals. *Ex vivo* engineered CAR T cells that recognise tumor cells are introduced into the patient, resulting in a living drug capable of a sensitive, target-specific, self-replicating, and long-term response [2, 3]. Despite success, further advancement of CAR T cell therapy is limited by many challenges including loss of persistence, T cell exhaustion and associated toxicities [4]. Additionally, patient responses vary in remission rates and associated toxicities such as cytokine release syndrome [5, 6]. To overcome these challenges, recent efforts have focused on designing novel CARs that show promising therapeutic characteristics, such as memory and cytotoxicity features and limited exhaustion [7]. For example, varying the signalling domains in the CAR was shown to lead to differences in T cell survival and cytotoxicity both *in vitro* and in patients [8, 9]. Still, the understanding of the exact properties of co-stimulatory domains that determine therapeutic efficacy is limited [10]. Despite efforts to establish high-throughput screening, current *in vitro* methods can profile 100 to 200 variants in primary (patient-derived) human T cells [11–14]. Although larger screens have been established, these rely on cell lines that do not completely recapitulate T cell biology [15]. However, currently up to three co-stimulatory domains can be incorporated in a CAR design, given the number of possible signaling domains involved in immunological function, this results in a CAR design space of over 100’000 possible combinations.

Computational modeling can help navigate this vast combinatorial space, mapping the relationship between CAR design and functionality and thus aiding lab-based screening by suggesting promising CAR T cell designs. For example, Daniels *et al*. recently trained a neural network model on a library of around 250 CAR variants to predict stemness and cytotoxicity from the CAR design for over 2000 unmeasured variants [13]. Although the authors were able to point to CAR design rules suggesting efficacy, they relied on summary statistics of the outcome that potentially mask heterogeneity of patient response. Indeed, during a CAR T cell therapy patients are infused with millions of cells, which can be best described by distributions rather than summary statistics [16, 17]. For example, one patient showed at the peak of the response 94% CAR T cells from a single clone, highlighting the importance of taking cellular heterogeneity into account [18]. Additionally, the authors used a one-hot encoding strategy to represent each CAR design as a combination of the thirteen signalling motifs assessed in their study, an approach that cannot generalise to other CAR T cell libraries.

Recently, the growing availability of single-cell RNA sequencing (scRNA-seq) data before and after perturbations has led to the emergence of machine learning models that predict gene expression distribution of single cells in response to drug or genetic perturbations, with early models based on different flavors of autoencoders [19–21]. As scRNA-seq is a cell destructive assay, an inherent difficulty in learning perturbation responses from the resulting data is that the same cell cannot be measured before and after a perturbation. As a result, control and perturbed cell populations are unpaired and direct cell-to-cell comparisons are impossible. Optimal transport (OT) is a field of mathematics concerned with moving mass between probability distributions in a cost-minimizing manner. It poses a natural framework to match the distributions before and after perturbations while accounting for the heterogeneity of the responses [22]. With neural OT, parametric models learn a global OT map through amortized optimization of before/after distributions which allows one to make predictions for unseen initial (cell) distributions. Neural OT has already been successfully employed in matching single-cell distributions over time [23–25], in space [26, 25], or across modalities [27– 29]. OT-based methods optimize the transportation map from a distribution of control cells to a distribution of perturbed cells. The resulting map can be used to infer a perturbation response, that is, to predict the change in cell state of the control cells if they had been exposed to this perturbation. The transportation can be learned for the (reduced) gene expression space or other modalities, such as cell surface markers [30]. Beyond the predictions of unseen cells [22], recent frameworks such as CondOT [31] and the conditional Monge Gap [32] allow to make predictions for unseen cells from unseen perturbations, thus extrapolating to new therapeutic options.

Inspired by the above, in this work we treat modeling of CAR T cell response as a perturbation prediction problem, and explicitly model response to a CAR T cell treatment at a single-cell level. We use OT as a generative model, which can generate the gene expression of a cell that expresses a CAR design that is not yet experimentally tested. We introduce a conditional CAR OT (CAROT) model, predicting the single cell distribution of thirty CAR variants. By leveraging recent advances in Protein Language Models (PLMs) that have successfully been used as embeddings for various downstream tasks [33], conditional CAROT (conCAROT) allows us to generalise to unseen CAR designs. We used the model to predict different CAR variants, obtaining distinct and accurate gene expression profiles that capture biological characteristics for in distribution CARs. Additionally, we made predictions for gene expression of unseen CAR T cells, showing that the conCAROT learned gene expression patterns linked to specific domains. Overall, we show that OT may aid discovery and development of new CAR T cell designs.

## 2 Methods

### 2.1 OT fundamentals and Conditional Monge Gap

OT is a mathematical framework that finds the optimal way to transport a source distribution to a target distribution while minimizing the cost of displacement. For a comprehensive overview of OT formalisms and biological applications, we refer to Bunne *et al*. [34]. The Monge formulation of OT [35] finds a push-forward map T that maps the source distribution to the target distribution while allowing splitting of the mass, and is formulated as

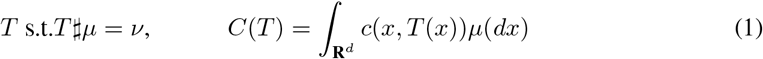

where µ and ν are measures on the source and target distribution, respectively, x is a point in the source distribution and *T* (*x*) the transportation of x. *T*#µ describes the result of transporting every point in µ and *C*(*T*) the cost associated with the transport. In single-cell applications, the Monge formulation translates to finding a map that transforms the gene expression of the control cells (source) to the gene expression of the perturbed cells (target). The mass splitting allows gene expression from a single source cell to be transported to multiple target cells, and a single target cell to receive gene expression from multiple source cells. In the case of neural OT, the learned transport map can then be applied to unseen cells to predict the target distribution at inference time. Such approaches often rely on Brenier’s theorem and use input convex neural networks (ICNNs) to parameterize and learn the OT maps [22, 36–39]. In practice, training ICNNs is challenging and can be circumvented e.g., with the Monge Gap [40], a regularizer that can be employed as a loss function in any neural network and estimates the deviation from a proposed transport map to an optimal map. The Monge Gap using n samples of a distribution is:

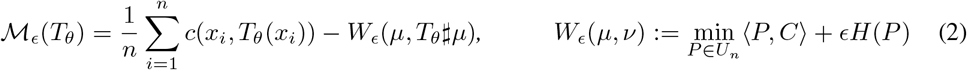

where c is again a cost function, *T*_θ_ a transport map, and µ and ν measures on the distribution (as in Equation 1), *W*_ϵ_ is the entropically regularised Kantorovich relaxation, with *H*(*P*) as entropy, ϵ the strength of regularisation, ⟨P, C⟩ the transportation cost using map *P* and cost matrix *C*, and *U*_*n*_ the set off all possible transport maps between the measures µ and ν. Here, we leverage the *conditional* Monge Gap (CMonge, [32]), a recently proposed extension of the Monge Gap that globally parametrizes Equation 2 by optimizing over *K* conditions simultaneously [32]:

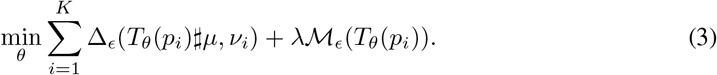

where ℳ _ϵ_ is the Monge Gap defined in Equation 2, Δ_ϵ_ is the Sinkhorn divergence between source and target samples [41], *p*_*i*_ describes the perturbation or condition, and λ is the strength of the Monge Gap regularization. This setting allows us to learn transport maps *conditioned* on a covariate and extend to unseen perturbations.

### 2.2 Conditional perturbation modeling of CAR T cell therapy

#### Chimeric Antigen Receptor library

We used a Chimeric Antigen Receptor (CAR) T cell library of 31 constructs, built from six intracellular signalling domains (Figure 1A) and cells of two donors from GEO database number GSE262686 [42]. The library consists of 30 CAR variant designs that contain either one (second generation) or two intracellular co-signalling domains (third generation) in addition to a CD3ζ domain. The control cells express a non-signalling CAR (NS-CAR) that only contains the antigen recognising extracellular domain specific for a cancer antigen, a trans-membrane domain, and no intracellular signalling domains domains. The cells underwent a repeated antigen stimulation study (RAS) to mimic *in vivo* conditions by continued T cell and tumor cell engagement and were subsequently sampled for scRNA-seq at day 0, 6, and 12 (Figure 1B). During the RAS, CAR T cells were co-cultured with a cancer cell line expressing an antigen that is recognised by all CAR variants for multiple days, to simulate exposure to cancer cells. The data were corrected for batch effects and annotated for CD4/CD8 cell states, cell cycle phase and functional scores. Functional scores were computed for cytotoxicity, memory, proinflammatory signature, T-helper 1 and T-helper 2 signatures using established marker genes [42] (subsection A.1, Table A1). This dataset is characterized by different sources of variation, including time, donor, cell type, and effect of CAR variant (Figure 1C). As expected, cells harvested at day 0 and days 6 and 12 formed distinct clusters, indicating large shifts in gene expression between short- and long-term activation. Another prominent source of variation is the cell state (phenotype): we observed seven CD8 cell states and eight CD4 cell states, including memory, activated, cytotoxic, bystander and dysfunctional states. The batch effect correction successfully eliminated confounding from donor variation. CAR variants were mixed in the UMAP embedding and only two CAR variants formed distinct clusters. The NS-CAR (brown) mostly occupied the resting memory compartments, showing a clear difference from cells that express CAR variants with signalling domains. The IL15R*α*-CD40-CD3ζ variant (turquoise) is prominent in the CD8 late bystander cluster.

**Figure 1:**
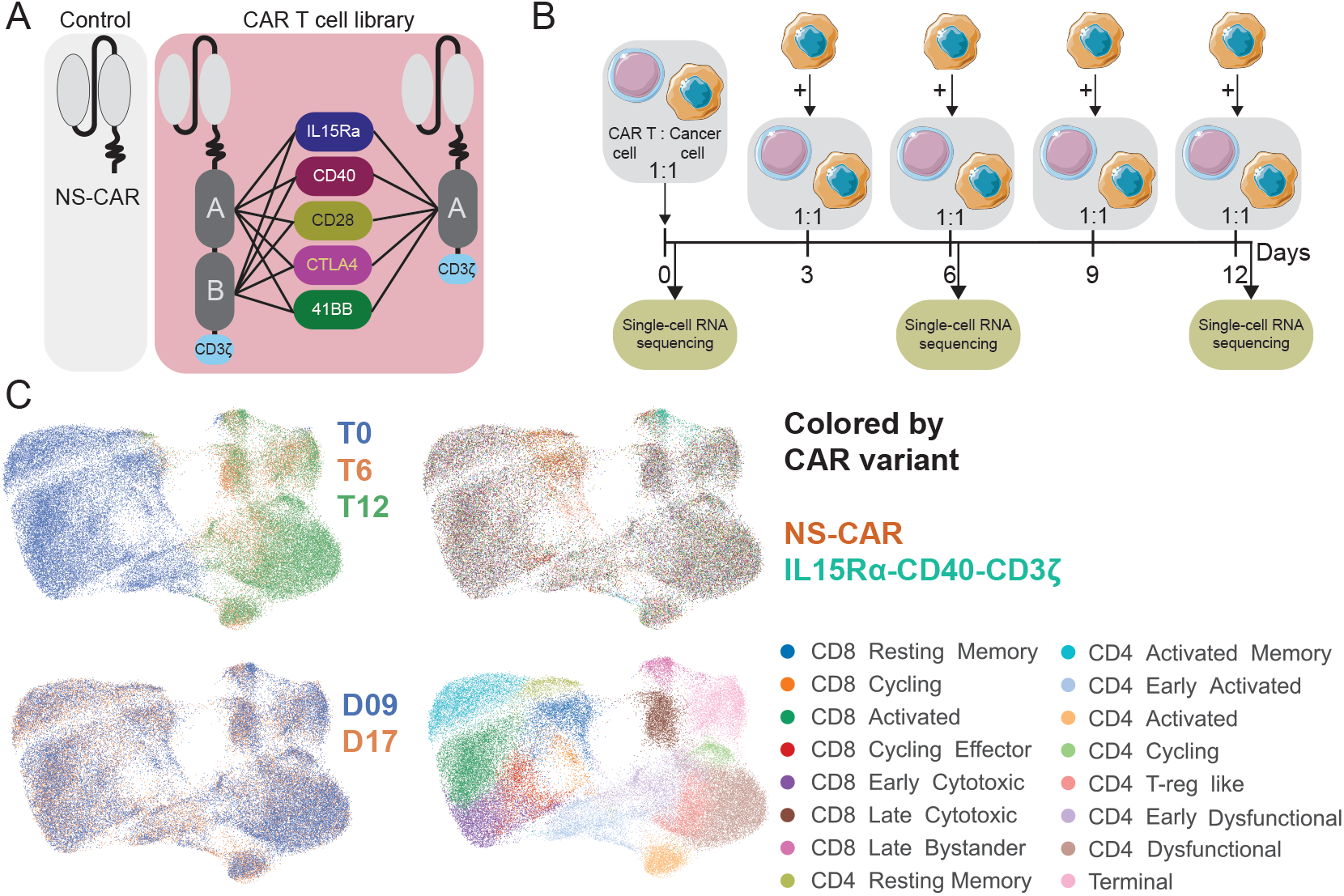
Dataset overview. A) CAR T cell library NS-CAR: Non-signalling CAR. B) Repeated antigen stimulation experiment setup. Cells for sequencing are sampled 6h after co-culture with the (newly added) cancer cells. C) UMAPs showing different covariates in the data based on the HARMONY representation of all genes.

#### CAR T cell OT

We build on the CMonge framework of (Harsanyi et al. [32], MIT license) and extend it to model response to CAR T cell therapy. We defined the NS-CAR cells as our source distribution and mapped it to target distributions of CAR-expressing cells (Figure 2A), transporting the non-batch corrected log-counts of 82 genes (see subsection A.2). We trained and applied CAROT on the CD4 or CD8 subsets separately, since transition from CD4 to CD8 states is rare and not the focus of this study. In contrast to previous works that used autoencoders to compress the scRNA-seq data [31, 22, 32, 20, 43], we used a curated functional geneset that contains 82 genes which capture T cell characteristics (e.g., memory, cytotoxicity, exhaustion and proliferation – Table A2), allowing for more interpretability (subsection A.2). We also investigated two baseline settings (Figure 2B): (i) an *identity* or *do-nothing* setting as the lower bound, as it takes cells from the control population without transportation as prediction for the CAR-expressing cells, and (ii) a *within-condition* setting that serves as an upper bound, as it takes CAR-expressing cells from the target CAR variant as prediction. Thus, ideally CAROT performs better than the identity model and close to the within condition model.

**Figure 2:**
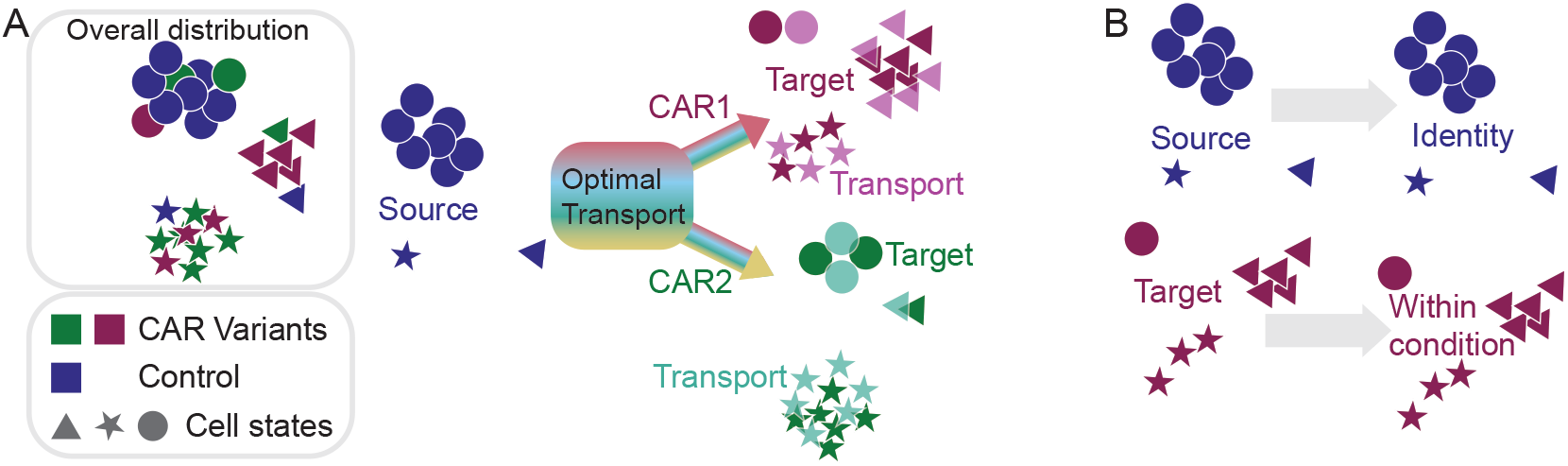
Optimal transport settings. A) The overall distribution shows control and CAR-expressing cells, clustered by cell state. Different variants have a different number of cells in the cell states. Optimal transport model CAROT maps gene expression of control cells (source) to gene expession of CAR predicting cells (target), trying to minimise the transportation cost and the error between the transport and the target expression. B) Two baseline settings. Identiy or do-nothing model uses gene expression of control cells and within condition takes gene expression of car expressing cells as prediction/transport.

We built on the CMonge implementation [32], which has condition-specific dataloaders for efficient GPU-use, meaning that each batch comes from a single condition/CAR variant. We mixed different conditions by using layer normalisation after concatenation of the embedding and gene-expression and take a gradient step after accumulation over four batches. For in distribution settings a train/test split of 0.8/0.2 was used. For out-of-distribution (OOD) experiments we used the same split for training, but, when evaluating the OOD variants, all cells were in the test set (0/1 split). We used the CMonge hyperparameters unless stated differently. We performed single-GPU training/evaluation with a runtime limit of 1h for the unconditional CAROT and 12h for the conCAROT models.

#### CAR embedding

To allow conCAROT to condition on a CAR variant and make CAR-specific predictions, we need to construct a generalizable embedding of the different CAR variants. Protein Language Models (PLMs) such as Evolutionary Scale Modeling (ESM), map the amino acid sequence to a numerical embedding and have been pretrained on different proteins and tasks, which allows them to capture biologically relevant information [44]. We embedded the intracellular amino acid sequence of the variants using ESM2 t48_15B_UR50D, referred to as ESM XL [44]. These embeddings were averaged over the sequence, resulting in the length of the embedding dimension of 5120 for ESM XL (subsection A.3).

## 3 Results

### 3.1 CAROT projects CAR effect on single cell distribution

We started with the simplest, non-conditional setting of training one OT model per CAR variant, referred to as CAROT (Figure 3A). Models were evaluated using the average *R*^2^ of the mean gene expression per gene over all cells as in [32, 22] (subsection A.5, Figure 3B) and the Maximum Mean Discrepancy (MMD), a distance metric between distributions that compares the gene expression on a finer scale than the mean expression (as in [22], subsection A.5, Figure 3C). For both metrics, OT significantly outperforms the identity model. In terms of R^2^ CAROT performs similar to the within condition setting. To evaluate the quality of the prediction, we visualized the source, target and transport cells onto a UMAP projection generated from the functional geneset’s non-batch corrected log-counts of all CD4 or CD8 cells (Figure A3). We show as an example two CAR variants, where we observe good mixing between target and transport cells, whereas the source distribution is distinct for the CAR-expressing cells and prediction (Figure 3D, see Figure A4&Figure A5 for all CARs). We also observed distinct transport patterns for different CAR variants, indicating that different transport plans are learned per CAR variant. CD40-CD40-CD3ζ has most cells in the cycling effector and early cytotoxic cell states, which is captured by CAROT. IL15R *α* -CD40-CD3ζ has most cells in the late bystander and late cytotoxic cell states, where also most of the transport is located. Taken together, these results establish that CAROT can be used to model CAR-variant effects.

**Figure 3:**
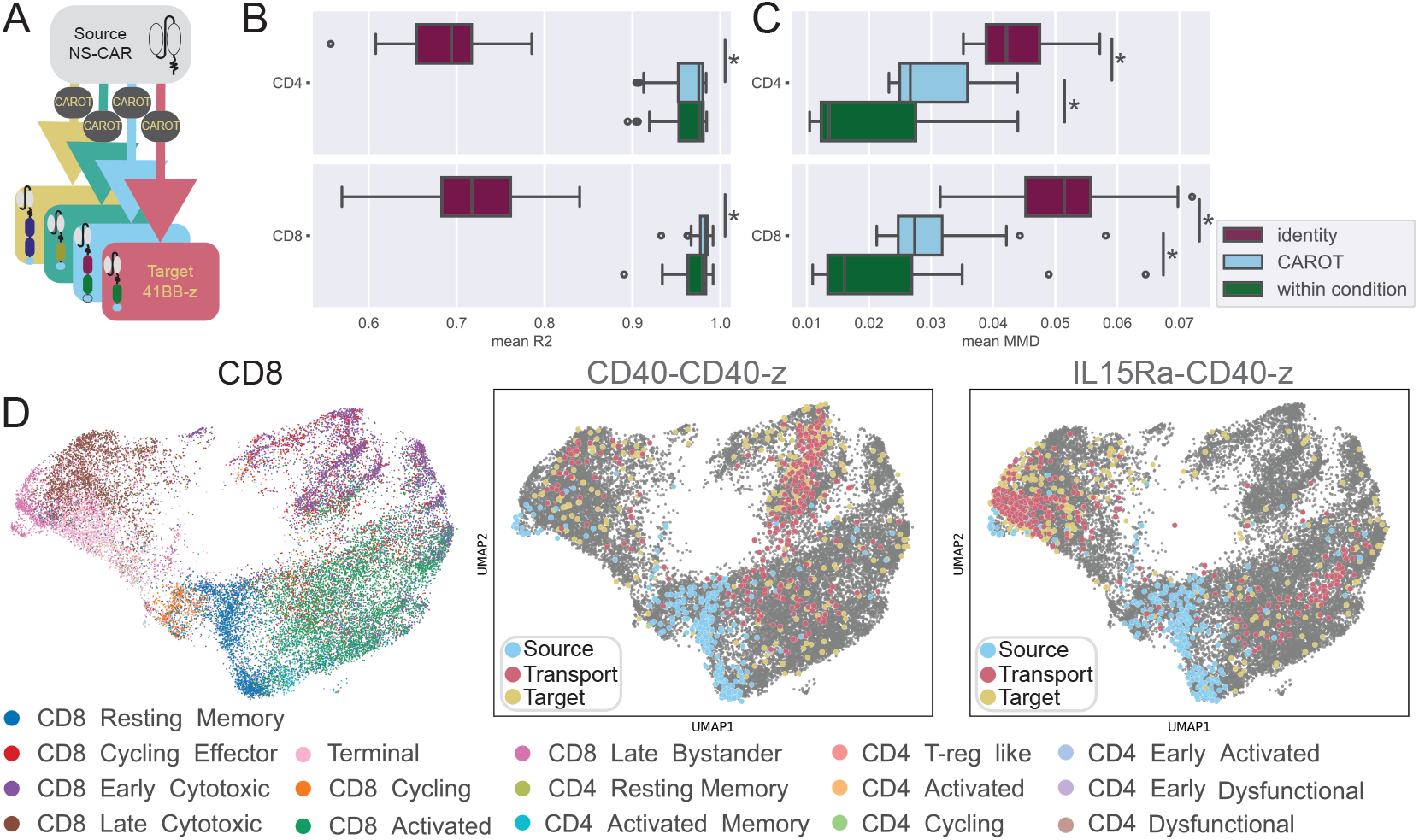
Unconditional optimal transport results. A) CAROT model setting. B) Mean *R*^2^ over 9 samples calculated using all 82 genes in the functional geneset for the CAROT model and the two baselines. Asterisks indicate statistically significant differences (subsection A.6). C) Same as B but showing mean Maximum Mean Discrepancy (MMD). D) UMAP constructed using the non-batch corrected logcounts of the 82 genes in the functional geneset for CD8 cells, colored by cell state (left), and with control cells (source), CAR-expressing cells (target) and prediction (transport) highlighted for two CARs (middle and right).

### 3.2 Conditional CAROT maps multiple CAR variants with one model

With the objective of predicting unseen CAR variants in mind, we trained a single OT model conditioned on a CAR variant embedding so that it can predict the effect of all CAR variants, we refer to this model as conditional CAROT (conCAROT) (Figure 4A). We observed that, for many CAR variants, the single cell distribution is quite similar, with few distinctions between CAR variants (Figure A4&Figure A5), which makes it hard for the model to learn to condition on the CAR variant. Also, for many CAR variants, there are only a small number of cells, complicating estimation of the single cell distribution (Figure A1B&C). Therefore, to train conCAROT, we selected only CAR variants for which we have more than 750 cells for CD4 and CD8 separately (Figure A1B&C).

**Figure 4:**
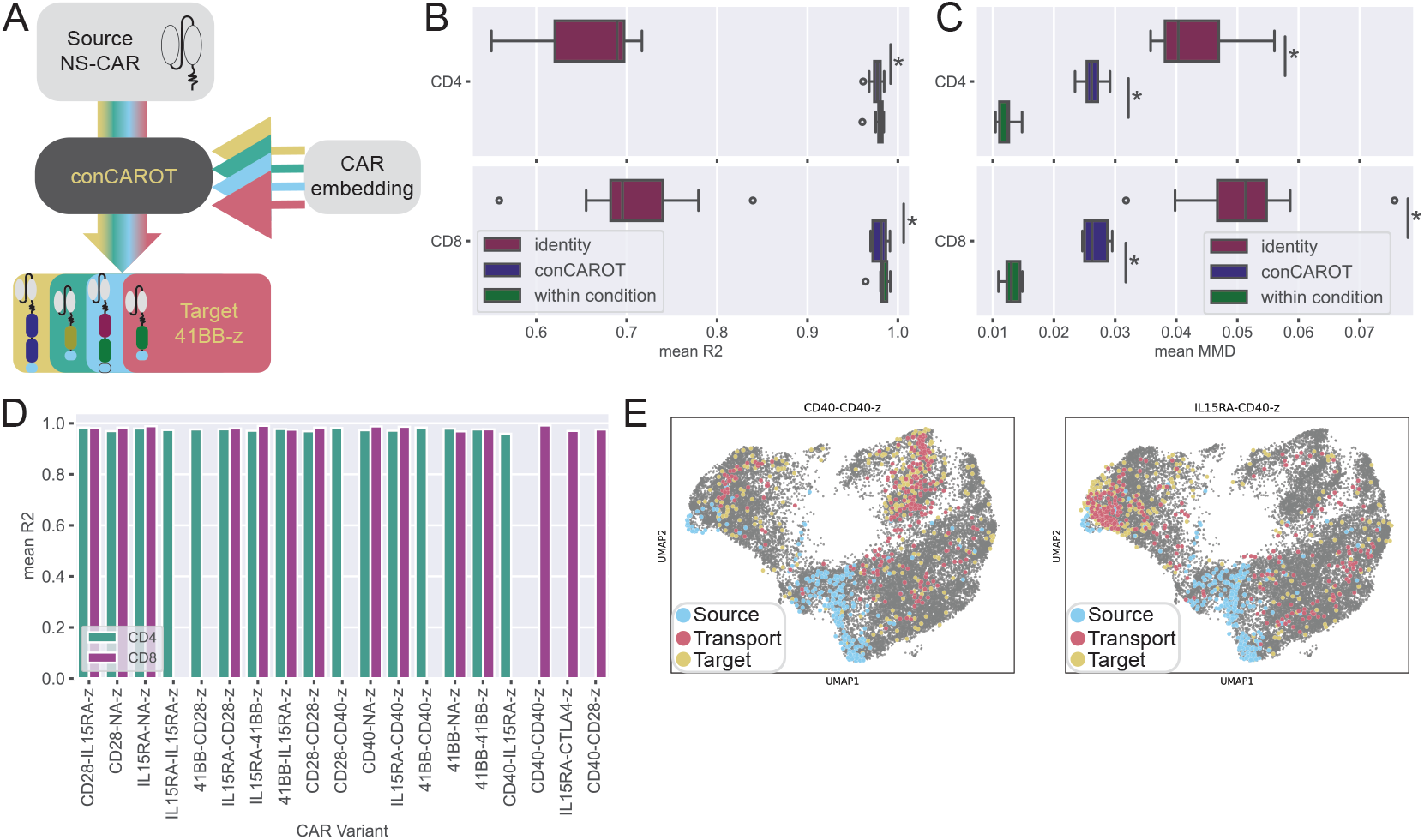
Conditional optimal transport results. A) Conditional CAROT model setting. B) Mean *R*^2^ over 9 samples calculated for the conCAROT model using the ESM XL embedding and the two baselines, and using only CARs with more than 750 cells. Asterisks indicate statistically significant differences (subsection A.6) C) Same as in B, but showing mean MMD. D) Mean R^2^ over 9 samples for CARs expressed in over 750 cells, split by CD4/CD8 subtype. Overall distribution is shown in panel B. E) Same UMAP as in Figure 3D (left), highlighted are the control cells (source), CAR -expressing cells (target) and prediction (transport) for two CARs.

We compared again the MMD and *R*^2^ with the identity and within condition models for both CD4 and CD8 (Figure 4B&C), achieving significant improvement over the baseline in both subsets and metrics. *R*^2^ values were greater than 0.95 for each CAR variant (Figure 4D) and comparable to the condition-specific CAROT models (Figure A10F), indicating no loss in performance compared to CAROT. As before, we see distinct transport patterns for different CAR variants, especially for the CD8 subset, where CD40-CD40-CD3ζ cells are found in the early cytotoxic and cycling effector cell states, and the IL15R*α*-CD40-CD3ζ cells found in the late bystander and late cytotoxic cluster (Figure 4E). The CD4 subset CAR-expressing cells differ less, although the predictions look well mixed with the target (Figure A6, CD8 Figure A7).

We further evaluated whether our predictions capture biological characteristics of CAR T cell therapy. We compared the distribution of functional scores, namely cytotoxicity, memory and proinflammatory scores, between the source, target and transport (Figure 5A). We observed similar medians and spread in the score distributions of cytotoxicity and memory of target and transport, unlike the ones between target and source. Furthermore, we compared the percentage of positive cells for each score and each CAR variant and marked that, for most variants, this showed good concordance between target and transport (Figure 5B), even for CAR variants that were not present in the training data (Figure 5B purple dots). However, the percentage of memory signature cells is often underestimated in the target for both in-distribution (ID) and OOD CAR variants, although the ranking of CARs based on the memory signature is maintained. This could indicate a poorer transport of memory genes or more difficult relation between CAR variant and memory signature, as CAR effect on the memory phenotype is less pronounced especially in the CD8 subset [42]. Moreover, we investigated the cell state distribution for different CAR variants and observe an overall similar pattern (Figure 5C).

**Figure 5:**
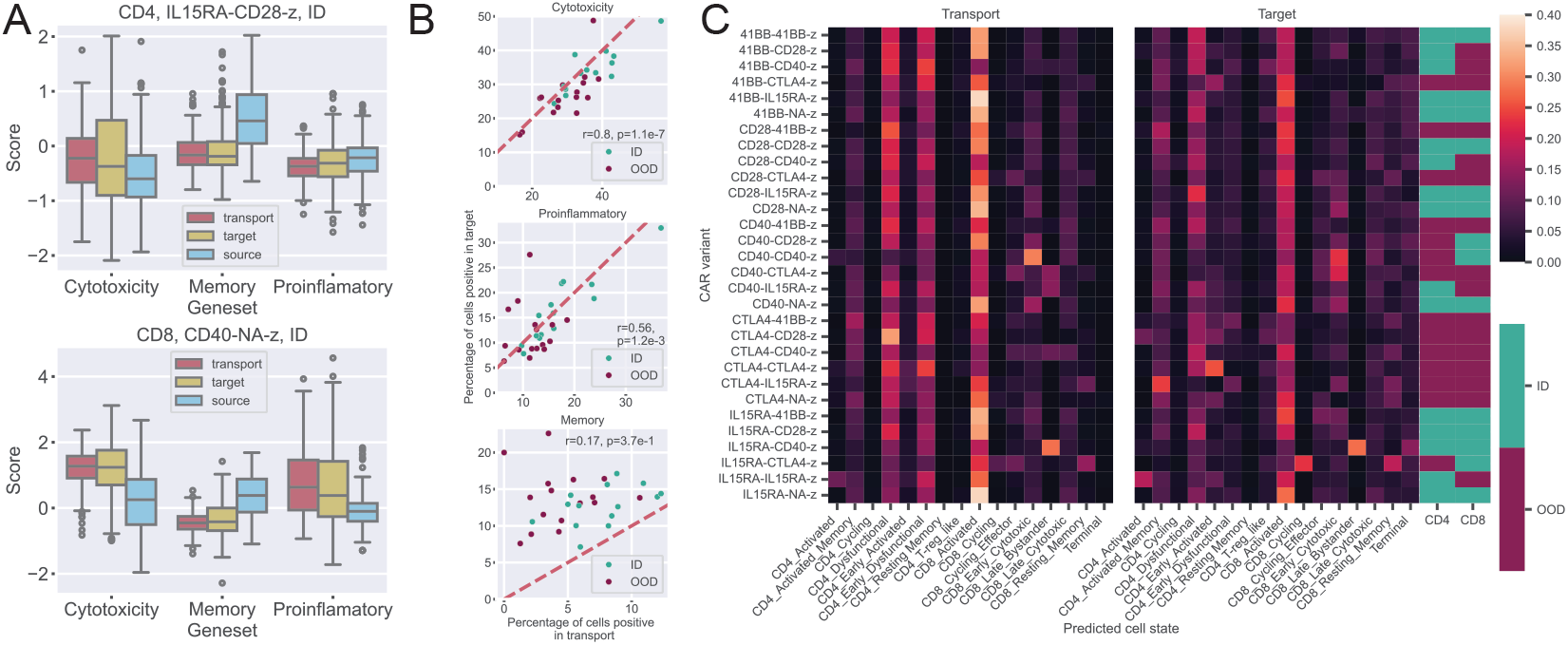
Biological features of transported gene expression. A) Geneset scores for two in-distribution (ID) CAR variants, colored by source, target and transport. B) Percentage of positive CD8 cells for each geneset in the target and transport, colored by ID and OOD variants. C) Heatmaps of predicted cell type fractions for transport and target, annotated with a label that indicates whether the CAR was ID or OOD for each subset.

conCAROT accurately captures the CD8 Early Cytotoxic cell state in the CD40-CD40-CD3ζ variant, the CD8 Late Bystander in the IL15R*α*-CD40-CD3ζ and the CD8 Resting Memory cell state in the IL15R*α*-CD40-CD3ζ variant. However, certain discrepancies between target and transport were also observed. For example, conCAROT tends to predict more cells in the CD4 Early Dysfunctional cell states than present in the target. Also, the CD4 Activated Memory and CD4 Early Activated populations in CTLA4 combined with IL15R*α* and CTLA4, respectively, are not well predicted by the model. This is possibly because we had to exclude most CAR variants with CTLA4 in position A due to low cell numbers. Overall, we conclude that conCAROT performs on par with CAROT and it manages to capture biological patterns, with the important benefit of a single model that makes predictions for multiple CAR variants.

### 3.3 Out-of-distribution prediction of CAR effects

Our goal is to have model that can make accurate predictions of unseen CAR variants, such that it enables *in silico* screening to select promising CAR variants for experimental testing (Figure 6A). Therefore, we further investigated conCAROT in the OOD setting (condCAROT-OOD). When we trained only on CARs with >750 cells, we automatically left out other variants with <750 cells. We observed that the OOD performance on the left out CARs still outperforms the identity setting significantly for the *R*^2^ but less explicit for the MMD (Figure 6B&C). Interestingly, the within condition and identity baselines also achieve a worse MMD on these held out CAR variants, suggesting that those variants are more challenging to predict, which is corroborated by further analysis (Figure A10A&B). Additionally, we observed that conCAROT-OOD transports many cells to the late cytotoxic or late bystander cell state when CD40 is present and many cells to the cycling or resting memory cell state for CTLA4 (Figure 6D). In the training data, the variants IL15R*α*-CD40-CD3ξ and IL15R*α*-CTLA4-CD3ξ clearly show clustering of cells in these regions as well, showing that conCAROT learns to condition on specific domains (All OOD CARs in Figure A8&Figure A9).

**Figure 6:**
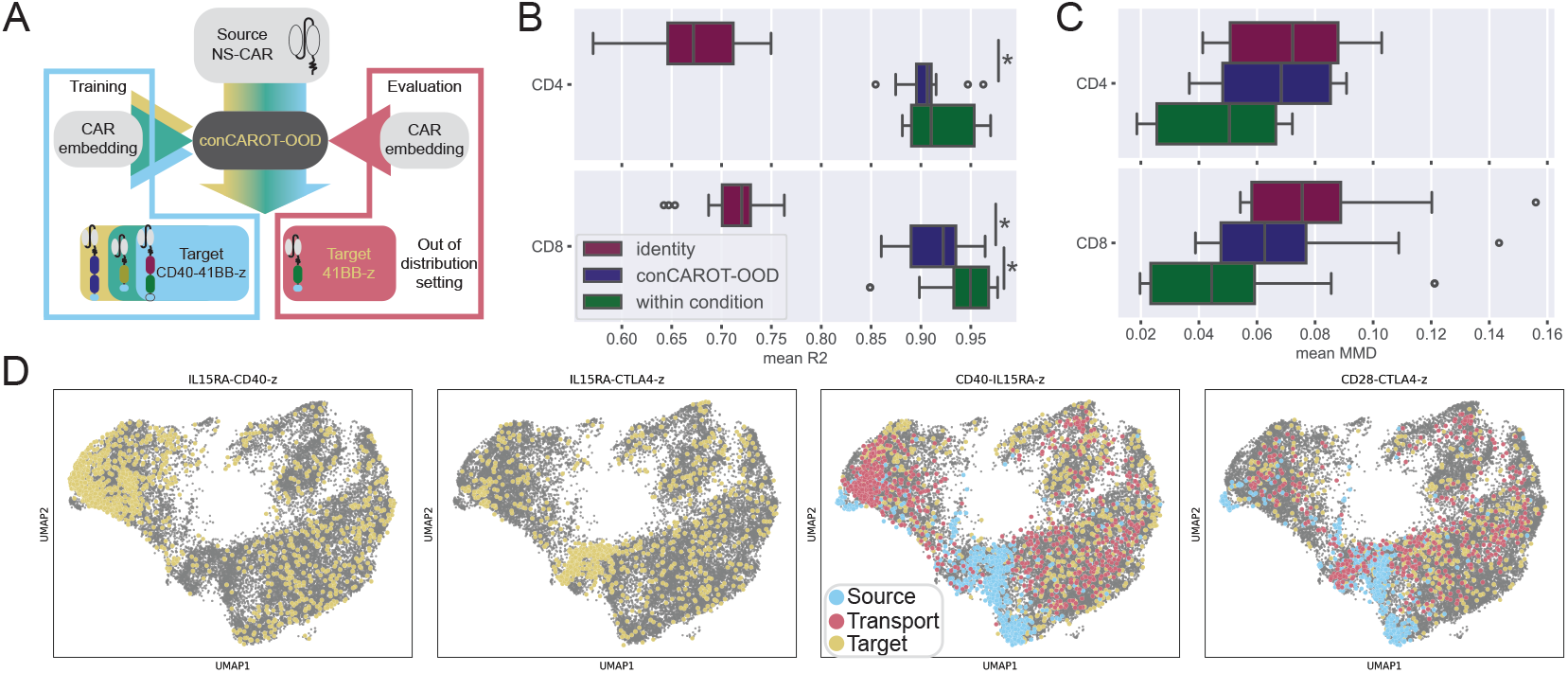
Out of distribution conditional optimal transport results. A) conCAROT OOD model setting. B) Mean *R*^2^ over 9 samples calculated for conCAROT and the two baselines using the ESM XL embedding for CARs not seen during training (CARs with <750 cells). Asterisks indicate statistically significant differences (subsection A.6). C) Same as in B, but showing the mean MMD. D) Same UMAPs as in Figure 3D, with all CD8 target cells for IL15R*α*-CD40-CD3ζ and IL15R*α*-CTLA4-CD3ζ shown in yellow in the first two subplots. These CAR variants were in the training data and show the CD40 and CTLA4 patterns that were learned. The third and fourth UMAPs highlight control cells (source), CAR-expressing cells (target) and prediction (transport) for two CARs, showing the OOD predictions for variants with CD40 or CTLA4.

To determine if conCAROT benefited from training on more CAR variants in the OOD setting, we trained 30 conCAROT models by leaving one CAR variant out of the training data and evaluating on the left out CAR variant. Again, conCAROT outperforms the identity model for both CD4 and CD8 in terms of the *R*^2^ and MMD, even on the OOD setting (Figure A10C&D). The UMAP projections for source, target and transport show good mixing of target and transport for CD4 CAR variants (Figure A11). For the CD8 subsets, we observed that it is difficult to capture the distinct patterns for different CAR variants, as variants with clear expression pattern show similar transport maps (Figure A12). This could again be due to many CAR variants having similar gene expression, hindering the model to condition on the CAR variant. However, the OOD setting is also harder and more CAR variants are likely needed to improve generalising to unseen CAR variants.

## 4 Limitations

A current limitation of this work is a lack of benchmarking to other perturbation prediction methods, which we are currently working on and will include in the future [21, 20]. Additionally, the results shown here are limited to one dataset based on healthy donors. Generalisation to other datasets or to CAR T cells used in cancer patients needs to be verified in future work. We would also like to point out that we did not specifically evaluate unseen domains. For the CD8 conCAROT-OOD models, all domains were always in the training data. However, for the CD4 conCAROT-OOD model trained only on CARs with >750 cells the CTLA4 domain was not observed at all during training. Nevertheless, we did not do an extensive analysis of this model’s performance on CTLA4 CARs specifically nor ran an experiment with deliberately held out domains. This setting is also planned in future work.

## 5 Discussion

We presented a novel and promising approach for modeling the effect of CAR design on the CAR-expressing T cell population that can exend to unseen CAR variants. Using optimal transport, a method which naturally handles unpaired measurements under multiple conditions, we map control cells with a non-signalling CAR to cells expressing different CAR variants. We showed that we can accurately infer perturbations of multiple CAR variants when conCAROT is conditioned on ESM embeddings of the CAR design. These predictions capture biological characteristics of CAR variants, making *in silico* CAR screening feasible. Additionally, we showed promising results of using the conCAROT to predict gene expressing for cells expressing unseen CAR variants. This model used domain-specific patterns learned from CARs in the training data to prediction for unseen variants.

This approach is adaptable to future indicators of clinical relevance and future CAR variants. Currently it is unclear which characteristics to use for *in vitro* CAR variant selection for clinical outcome [45]; many possible genesets and downstream analyses are possible. A future study could point to new clinically relevant indicators that can be derived from gene expression, not just the functionalities or cell states investigated here. Different downstream analyses on the predicted gene expression could facilitate easy adaption to novel CAR T cell characteristics. Additionally, we could also retrain the model using novel genes if the novel indicators are not well captured in the current geneset. As we use a protein language model for embedding the CAR variants, all future CAR variants can be encoded as long as they can be represented with an amino acid sequence. Previous CAR T cell models focused only on stemness and cytotoxicity and encoded variants with a binary encoding [13]. This prevents the previous models from extending to CARs with a different library design and adaptation to other readouts without retraining.

Increasing CAR library size could improve OOD generalisation, especially when it includes more CARs with distinct patterns. We observed that many CAR variants responded similarly in the repeated antigen stimulation setting. This hinders the model’s ability to condition on the CAR embedding, since the variable that it is conditioned on can have little effect. Together, this hampers the accurate prediction of CAR variants with distinct effect. A larger library could overcome some of these hurdles, as it gives the model more examples from which to learn more general patterns. Also, a larger library would most likely have more variants with distinct effects, improving the model’s prediction of distinct CAR variants.

A CAR-specific model could help screen patients for approved CAR T cell therapies. Patients have different responses and success rates for given (approved) CAR T cell therapies [2]. A model specialized on making predictions for one CAR variant from different control populations could make valuable predictions given a simple blood sample from patients. These predictions could give indication whether or not the patients might have a promising T cell phenotype distribution given the CAR variant, for example, indications of memory response or cytotoxicity of the resulting CAR T cells. In this study, we did not have enough data to investigate this approach, as splitting by donor leads to too few cells to effectively train the optimal transport model.

## 6 Data and code availability

Code is available under https://github.com/AI4SCR/CAR-conditional-monge/ and data will be accessible upon publication of the data paper under in the GEO database under accession number GSE262686 [42].

## 7 Acknowledgements

This project received funding from the European Union’s Horizon 2020 research and innovation program under the Marie Skłodowska-Curie grant agreement no. 955321, IBM Research and NCCR Molecular Systems Engineering.

