## Appendix for "Modeling CAR Response at the Single-Cell Level Using Conditional Optimal Transport"

### A Appendix

#### A.1 scRNA-seq preprocessing

The scRNA-seq data was quality controlled with nFeature_RNA > 300, nCount_RNA < 50000, percent.mt < 20 and nCount_ADT < 30000. Because of large differences in cell numbers per CAR variant, we applied a subsampling strategy. If a CAR variant has more than 6.25% of the cells in a sample, then these cells downsampled to maximum 6.25% of all cells. (6.25% is deduced from twice the average number of cells per variant 100/32 =3.123%). Cells were scaled and regression was done for cell cycle phase (S and G2M) and percentage of mitochondrial genes. Batch correction for donor and timepoint was done with Harmony, using lambda=c(1,200) for UMAPs in Figure 1 only [46]. This preprocessing was done before downloading the data and cell numbers per variant after quality control and sampling are shown in Figure A1. For detailed experimental set up and scRNA-seq preprocessing see [42].

#### A.2 Genesets and geneset scoring

The functional geneset was obtained by selecting 77 functional genes from various published research with scRNA-seq of CAR T cells and listed in Table A2. We included five additional genes needed to compute the functional scores listed in Table A1, resulting in the functional geneset of 82 genes. 82 random genes were sampled from genes that are expressed by at least one cell in each CAR variant. The highly variable genes (HVG) were determined by first taking the top 100 HVG of each CAR variant with the control, yielding 30 HVG genesets of 100 genes. Then we counted how often each gene occurred over all the 30 genesets and took the genes that occurred at least 22 times, giving us 81 unique genes. The number 22 was chosen to get a geneset size in the same range as the functional and score geneset size. We decided the final geneset based on the performance of unconditional CAROT and used this geneset for the conCAROT experiments.

To evaluate if the geneset captured differences between control and CAR-expressing cells, we calculated, for each gene, its information content in terms of entropy between CAR-expressing cells and control cells (Figure A2) and compared it with the one obtained from 82 randomly sampled genes and 81 HVG. We observed that the functional and HVG genesets scored similarly, indicating that our functional geneset can differentiate between control and CAR populations equally well as the HVG. To compute the information content of each gene across control and CAR treatment, we computed the relative entropy of the distribution of the sum of library-size normalised counts as follows

$$D = \sum_{k=0}^{G} (p_k * log(p_k/q_k))$$

where $p_k$ and $q_k$ are the counts of a geneset (of length $G$) for the control and the CAR-expressing cells respectively, normalized such that $\sum p_k = \sum q_k = 1$ [47].

**Table A1:** Genes used for geneset scoring. Taken from [42]

| Cytotoxicity | Proinflamatory | Memory | CD4_Th1 | CD4_Th2 |
|---|---|---|---|---|
| GZMB | IFNG | TCF7 | IL2 | IL5 |
| PRF1 | TNF | SELL | IFNG | IL13 |
| FASLG | CRTAM | CCR7 | TNF | IL4 |
| | CSF2 | LEF1 | | |
| | XCL1 | IL7R | | |
| | XCL2 | | | |
| | CCL1 | | | |
| | CCL4 | | | |

#### A.3 CAR embeddings

To construct the CAR embeddings, we compared different approaches to encode the CAR variant: (i) two binary embeddings, (ii) two ESM-based embeddings, and (iii) a metadata embedding. The two

**Table A2:** Prior knowledge functional genes from literature. CRS: Cytokine Release Syndrome

| Function | Gene | Refs | Function | Gene | Refs |
|---|---|---|---|---|---|
| Memory | CD3E | [48] | Exhaustion | ADORA2A | [48] |
| Memory | CCR7 | [49, 48] | Exhaustion | BATF | [50, 48, 51, 49] |
| Memory | CD28 | [48] | Exhaustion | BATF3 | [51] |
| Memory | CD27 | [48] | Exhaustion | BTLA | [48, 51, 52, 49] |
| Memory | SELL | [49, 48] | Exhaustion | CCL1 | [49] |
| Memory | IL7R | [53] | Exhaustion | CCL3 | [49, 52] |
| Memory | TCF7 | [54] | Exhaustion | CCL4 | [49] |
| Memory | LEF1 | [54] | Exhaustion | CCL5 | [49] |
| Memory | KLF2 | [53] | Exhaustion | CD160 | [55] |
| Cytotoxicity | GNLY | [56, 50] | Exhaustion | CD2 | [48] |
| Cytotoxicity | GZMK | [56, 52] | Exhaustion | CD244 | [52, 49, 55] |
| Cytotoxicity | GZMA | [50, 52] | Exhaustion | CD3E | [48] |
| Cytotoxicity | GZMB | [50, 52] | Exhaustion | CTLA4 | [55] |
| Cytotoxicity | PRF1 | [52] | Exhaustion | ENTPD1 | [49] |
| Cytotoxicity | LAG3 | [52] | Exhaustion | GZMB | [51] |
| Cytotoxicity | NKG7 | [52] | Exhaustion | HAVCR1 | [48] |
| Cytotoxicity | ZEB2 | [56, 54] | Exhaustion | HAVCR2 | [48, 52] |
| Cytotoxicity | EOMES | [56] | Exhaustion | ID2 | [50, 49] |
| Cytotoxicity | ZNF683 | [56] | Exhaustion | IFNG | [51] |
| Cytotoxicity | TBX21 | [54] | Exhaustion | IL13 | [51] |
| Cytotoxicity | PRDM1 | [54] | Exhaustion | IL17RA | [51] |
| Proliferation | IL2 | [56, 48, 53] | Exhaustion | IL2RA | [51] |
| Proliferation | LIF | [56] | Exhaustion | IRF4 | [51] |
| Proliferation | CENPV | [56] | Exhaustion | KIR3DL1 | [48] |
| Proliferation | G0S2 | [56] | Exhaustion | KLF2 | [51] |
| Proliferation | ORC6 | [56] | Exhaustion | KLRG1 | [52] |
| Proliferation | CD3E | [48] | Exhaustion | LAG3 | [55] |
| Proliferation | CD2 | [48] | Exhaustion | LAYN | [55] |
| Proliferation | CD28 | [48] | Exhaustion | LEF1 | [51, 49] |
| Proliferation | IL2RA | [48, 51] | Exhaustion | NCAM1 | [48] |
| Proliferation | CD69 | [48] | Exhaustion | NCR1 | [48] |
| Proliferation | ICOS | [48] | Exhaustion | PDCD1 | [50, 55] |
| Proliferation | TNFRSF4 | [48] | Exhaustion | TCF7 | [51] |
| Proliferation | TNFRSF9 | [48] | Exhaustion | TIGIT | [52] |
| Proliferation | CD27 | [48] | Exhaustion | TNFRSF18 | [51] |
| Proliferation | TNF | [48, 53] | CRS | IL1B | [50] |
| Proliferation | IFNG | [48, 51, 53] | CRS | CXCL8 | [50] |
| Proliferation | GZMB | [51, 53] | CRS | CCL3 | [50] |
| Proliferation | MKI67 | [52] | CRS | CCL4 | [50] |
| Proliferation | CDK1 | [52] | CRS | IL13 | [50] |
| Proliferation | CCNA2 | [52] | CRS | CD69 | [50] |
| Proliferation | CDCA2 | [52] | CRS | LEF1 | [50] |
| Proliferation | FOS | [49] | CRS | IL7R | [50] |
| Proliferation | CCL3 | [53] | CRS | STAT1 | [50] |
| Proliferation | CCL4 | [53] | CRS | FOXP1 | [50] |
| Proliferation | NCR1 | [48] | CRS | CD27 | [50] |
| Proliferation | NCAM1 | [48] | CRS | IL16 | [50] |
| Cytokines | CCL3 | [55] | CRS | GZMB | [50] |
| Cytokines | CCL4 | [55] | CRS | GZMA | [50] |
| Cytokines | CCL20 | [55] | CRS | BATF | [50] |
| Cytokines | IFNG | [55] | CRS | GZMH | [50] |
| Cytokines | IL10 | [55] | CRS | IL13 | [51] |
| Cytokines | TNF | [55] | CRS | IL1A | [51] |
| Cytokines | LAG3 | [55] | CRS | CSF2 | [51] |
| Cytokines | CD226 | [55] | | | |
| Cytokines | HAVCR2 | [55] | | | |
| Cytokines | HOPX | [55] | | | |

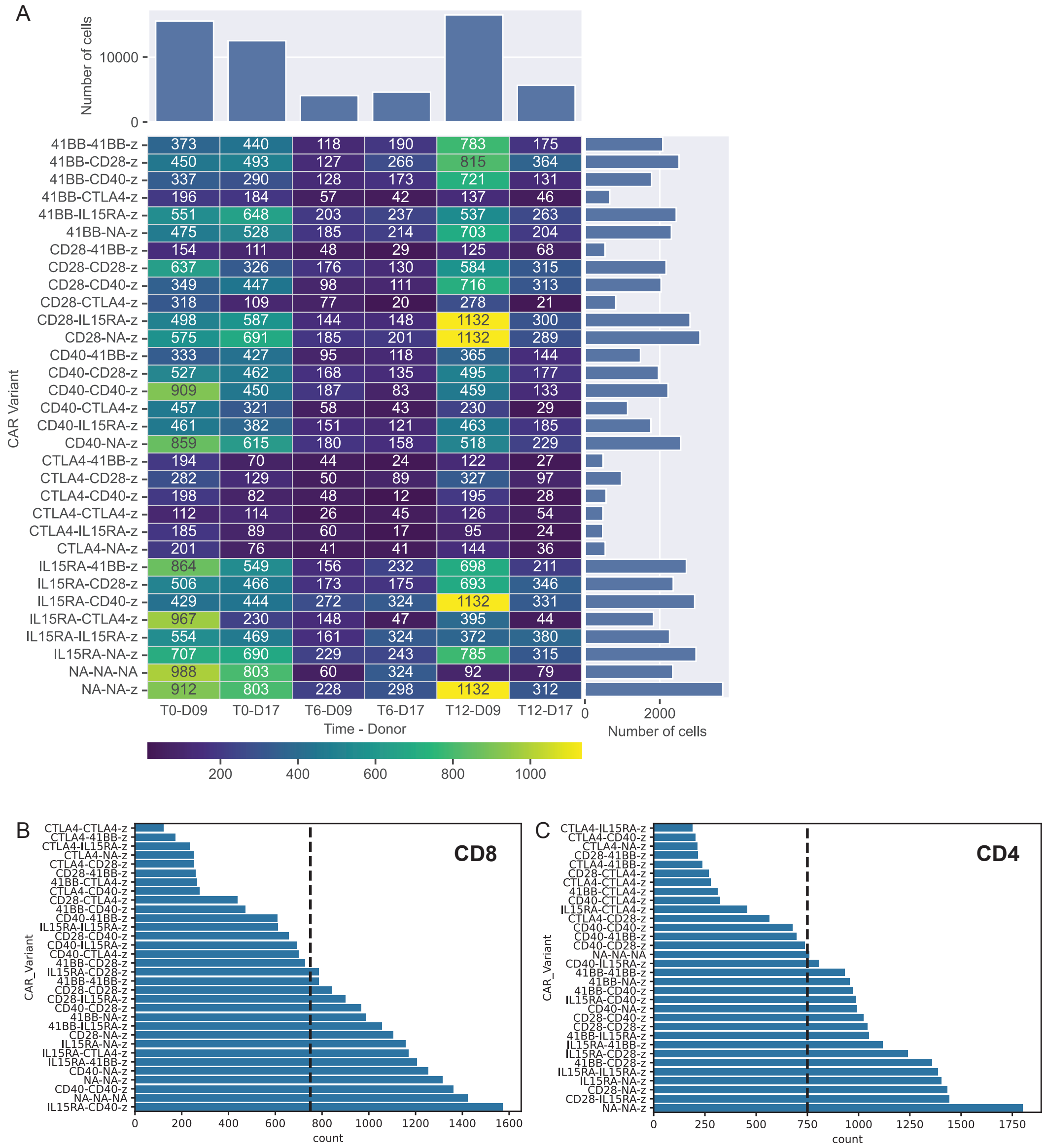


**Figure A1:** Dataset overview and CAR selection thresholds. A) Overview over cell counts per CAR variant, donor and timpoint. Cells are colored and annotated with cell counts, barplots on top and right show the marginal counts. B) Cell counts per CAR variant in the CD8 subset. Dashed line indicates threshold for selecting the variants for training data. C) Cell counts per variant in the CD4 subset. Dashed line indicates threshold for selecting the variants for training data.

binary CAR embeddings are based on the absence or presence of the five signalling domains and the CD3$\zeta$ domain. For the 11-dimensional embedding, the first five bits describe which domain is in position A. Each position corresponds to one of the five possible domains, and of these five bits only the bit corresponding to the domain will have a 1. Then there are five bits that in a similar manner describe which domain is in position B. The last and 11th bit describes the presence (1) or absence (0) of the CD3$\zeta$ domain. Similarly, the last bit of the 16-dimensional binary embedding also indicates CD3$\zeta$ presence. The 16d embedding has three bits for each of the five signalling domains. Of these three bits, the first position is 1 if the domain is at all present in the CAR variant, irrespective of position. The second bit describes whether the domain is in position A and the third bit position B.

We used Facebook's ESM2 models, either t6_8M_UR50D or t48_15B_UR50D, referred to as ESM small or ESM XL respectively to compare large and small ESM embeddings [44]. We embedded only the intracellular signalling tail, since this is where the CAR variants differ. Then we averaged these embeddings over the sequence, resulting in the length of the embedding dimension of 5120 for ESM XL and 256 for ESM small.

Additionally we used a metadata embedding, which includes information cells that express the CAR variant. For each variant the embedding consists of the means and standard deviation of the scores "Cytotoxicity_1", "Proinflamatory_2", "Memory_3", "CD4_Th1_4", "CD4_Th_5", "S.Score", "G2M.Score". Also, the fraction of cells from each of the donors, timepoints, cell cycle phases and cell states were included, resulting in a 40 dimensional embedding.

The binary embeddings are based on the CAR library used here and cannot embed novel CAR variants, whereas the ESM embeddings are based on the amino acid sequence of the variants and can therefore be extended to novel CAR variants. The metadata embedding captures considerable information that is derived from the scRNA-seq and should therefore greatly facilitate the conditional OT problem. We observed that a larger ESM embedding (ESM XL) performs better than a smaller ESM model (Figure A2). Interestingly, the binary embeddings don't show significantly worse performance than the metadata or ESM embeddings, despite having a lower dimension and only information about the presence/absence of domains. We continued with the ESM XL embedding, since it can be used in an out-of-distribution setting.

##### A.4 OT UMAPs and biological scores

For comparing biological scores between source, target and transport we sampled the same number of cells from source and target. The number of cells is the minimum of number of cells in the target or source validation set, as the target validation set is CAR variant dependent. Then we transported the target cells to obtain the same number of transport cells. All three datatypes (source, target and transport) were then mapped onto a UMAP based on all CD4 or CD8 cells.

Additionally, we calculated geneset scores for the cytotoxicity, memory and proinflammatory genesets. The thresholding for cells positive for a certain scores are taken from [42] and are 0 for the memory geneset and 1 otherwise. We trained a cell-typing model on all CD4 and CD8 cells, since this seemed to work better dan CD4 and CD8 separately. This Support Vector Machine model was then used to predict the celltypes for source, target and transport. This model was implemented with scikit-learn [57].

##### A.5 Metrics and evaluation

We leveraged the coefficient of determination ($R^2$) and the Maximum Mean Discrapancy (MMD) for model evaluation, as these metrics are often used in perturbation single cell models and optimal transport models [22, 31, 32, 20, 58].

For the $R^2$ the average expression per gene over all cells is computed, for the prediction and target separately. Then the R2 is determined on the average expression of all genes.

$$R2 = 1 - \frac{\sum_{i=1}^{g}(x_i - y_i)^2}{\sum_{i=1}^{g}(x_i - \bar{x})^2}$$

Where $g$ indicated all genes, and $x_i$ is the average target gene expression for gene $i$ and $y_i$ the average predicted gene expression for gene $i$. $\bar{x}$ is the average gene target gene expression over all genes.

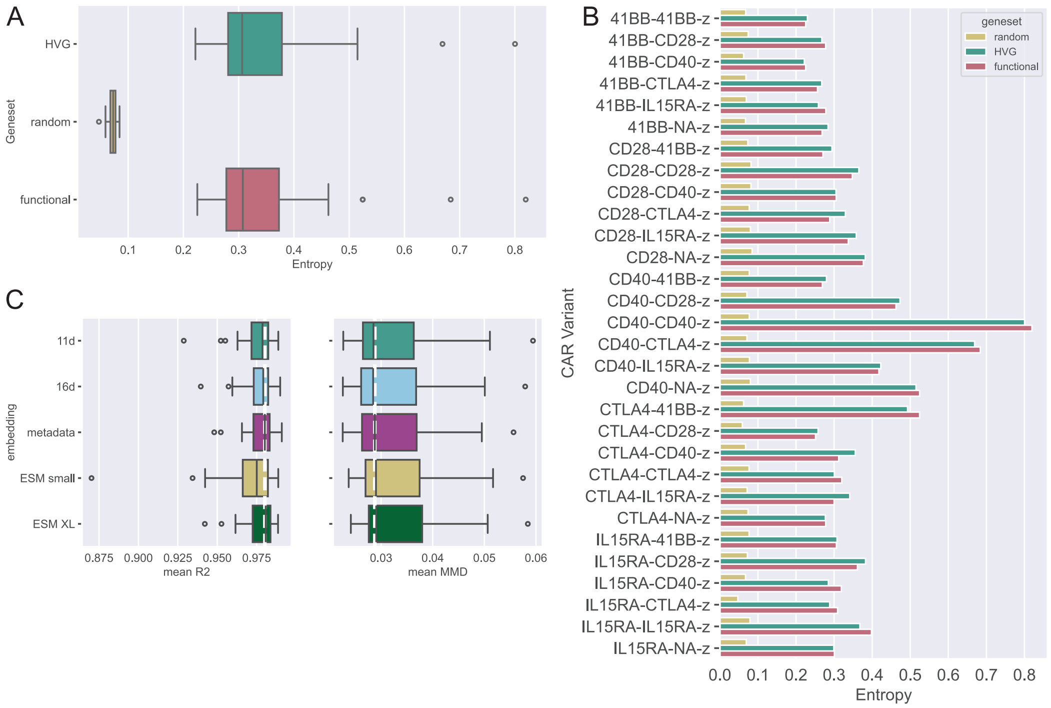


**Figure A2:** Genesets and CAR embedding evaluation. A) Distribution of entropy between the control gene expression counts and CAR-expressing cells gene expression counts, for genes in the indicated genesets. B) Entropy between control and CAR-expressing cells as in A, only now shown per CAR. C) Comparing different CAR embeddings using the conditional CAROT model, trained on all CAR variants and makes predictions for all CAR variants using a test split. The white line shows the median of the metadata embedding. Left shows the R2 and right the MMD.

We used the MMD to find the maximum distance between these distributions in the kernel space.

$$MMD(T, P) = \|\mu_T - \mu_P\|_K$$

where $T$ is the target distribution over all cells and genes and $P$ the predicted distribution. We take an average over the MMD with a RBF-kernel with $\gamma = [2, 1, 0.5, 0.1, 0.01, 0.005]$.

##### A.6 Statistical analyses

We tested (con)CAROT(-OOD) versus identity and (con)CAROT(-OOD) versus the within condition using a two-sided Mann-Whitney U test using [47]. This non-parametric test assumes independence between the two groups and ordinal observations. The independence assumption might not hold for our data, since the target difficulty and subsequent score might influence the performance of (con)CAROT(-OOD). We did multiple hypothesis correction using the Bonferroni method, by lowering the significance threshold $\alpha = 0.05$ to $\alpha = 0.05/n$ with $n = 8$ since we tested for each experiment two subsets, two scores and two comparisons ($2^3 = 8$). Significant differences are indicated with an '*' in the manuscript.

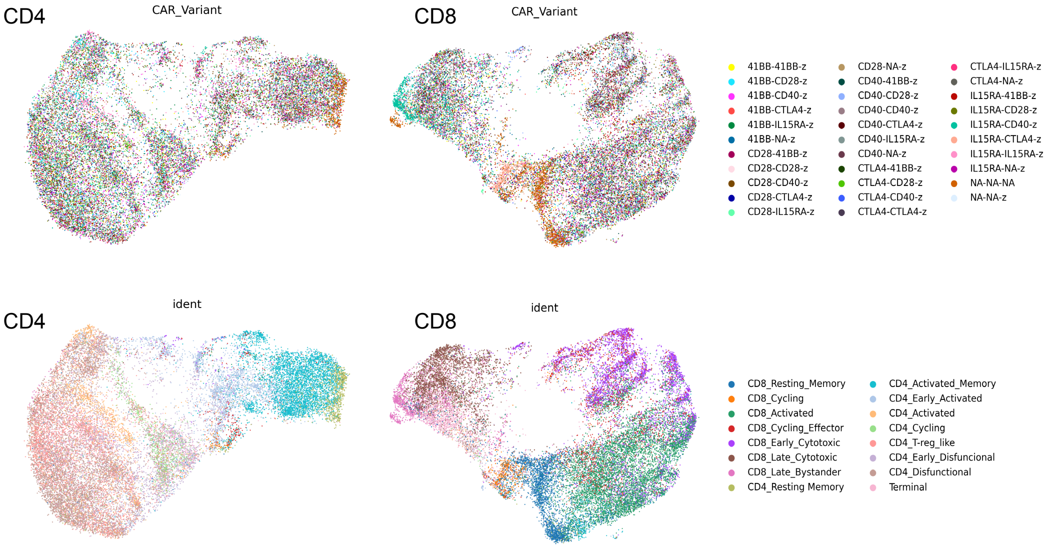


**Figure A3:** UMAPs of CD8 and CD4 subsets based on the logcounts of the 82 genes in the functional geneset. Colored by CAR variant and cell state.

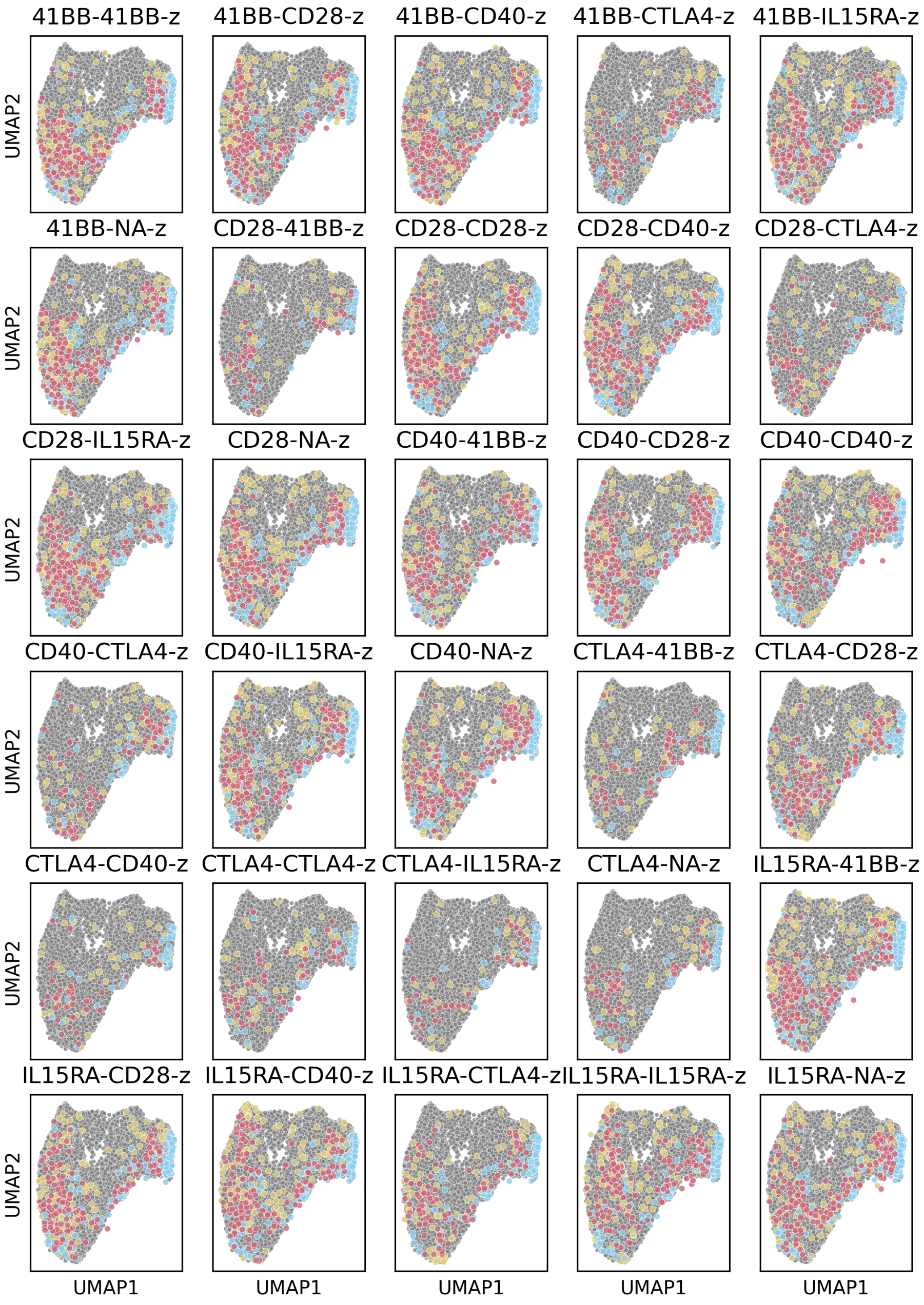


**Figure A4:** UMAPs of CD4 subset cells for the unconditional OT model (one model per CAR variant and subset).

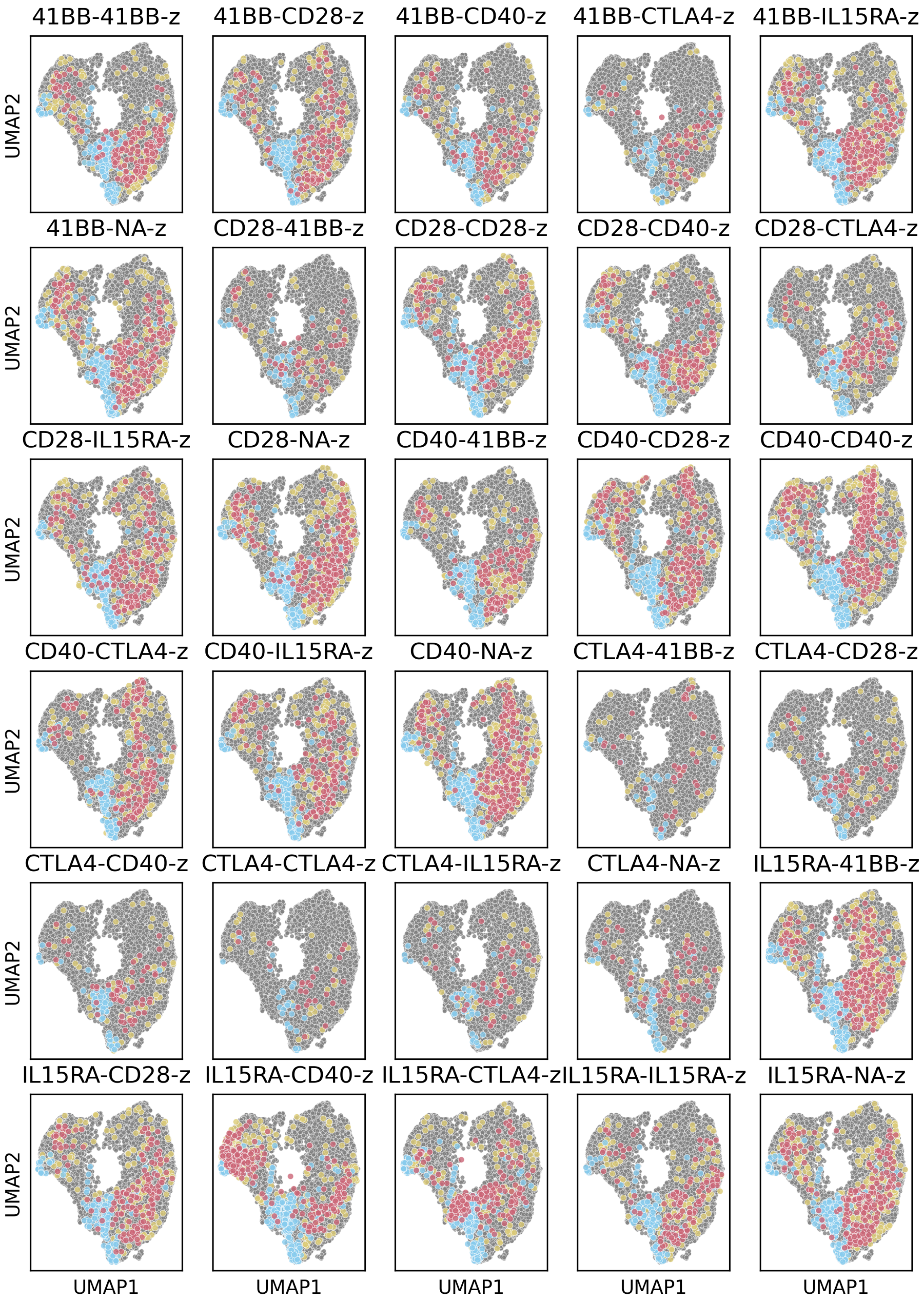


**Figure A5:** UMAPs of CD8 subset cells for the unconditional OT model (one model per CAR variant and subset).

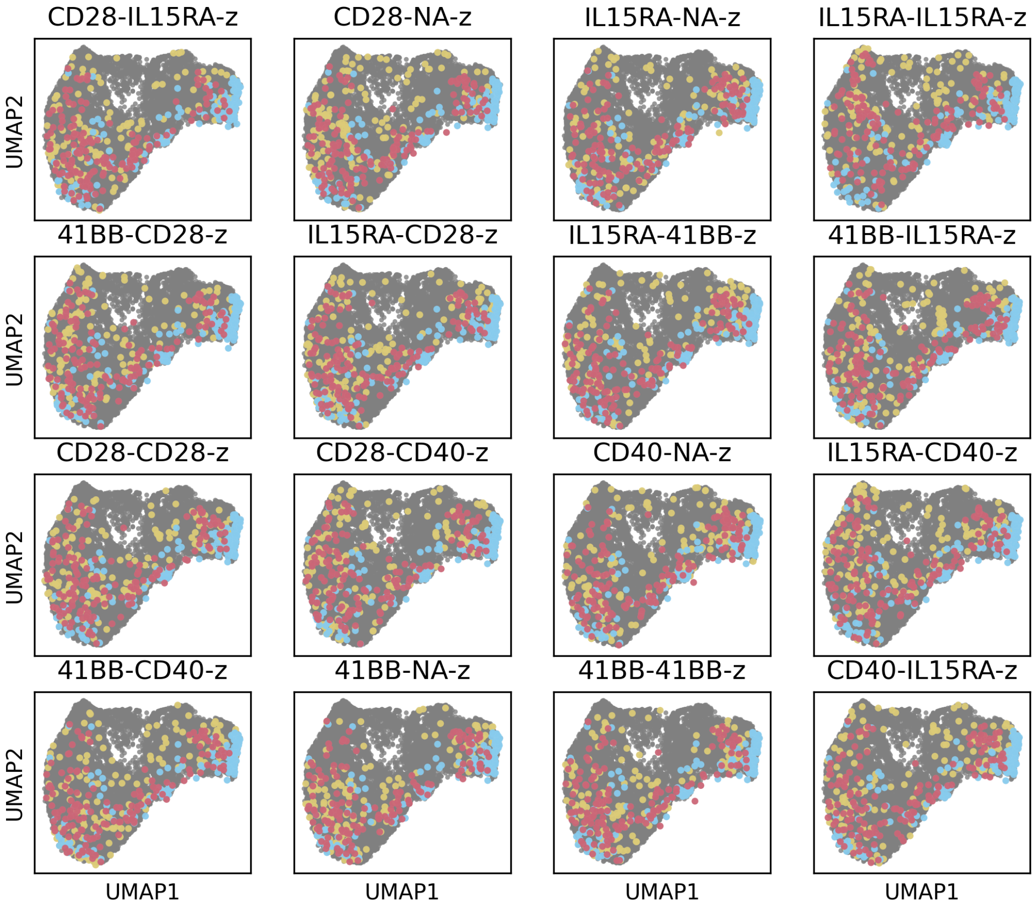


**Figure A6:** UMAPs of CD4 subset cells for the conditional OT model for CAR variants with >750 cells in the subset, all variants shown here were present on in the training set. Evaluation on a held-out test set.

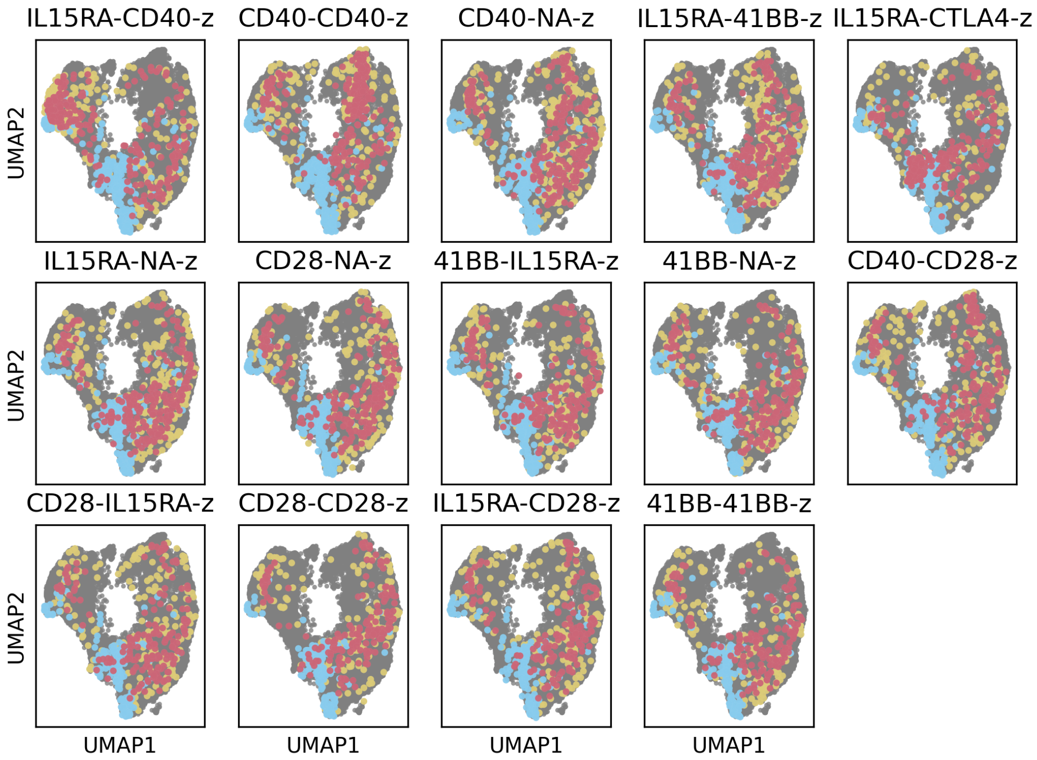


**Figure A7:** UMAPs of CD8 subset cells for the conditional OT model for CAR variants with >750 cells in the subset, all variants shown here were present on in the training set. Evaluation on a held-out test set.

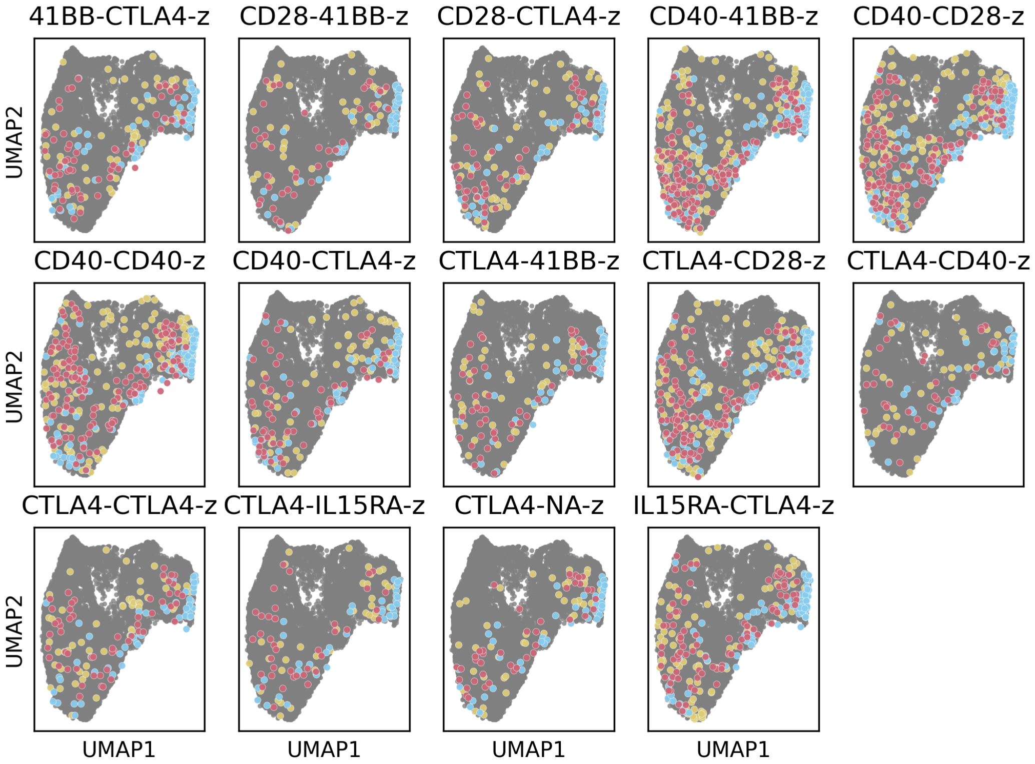


**Figure A8:** UMAPs of CD4 subset cells for the conditional OT model trained on CAR variants with >750 cells in the subset. All variants shown here were not present on in the training set, shown are all available cells for the variant

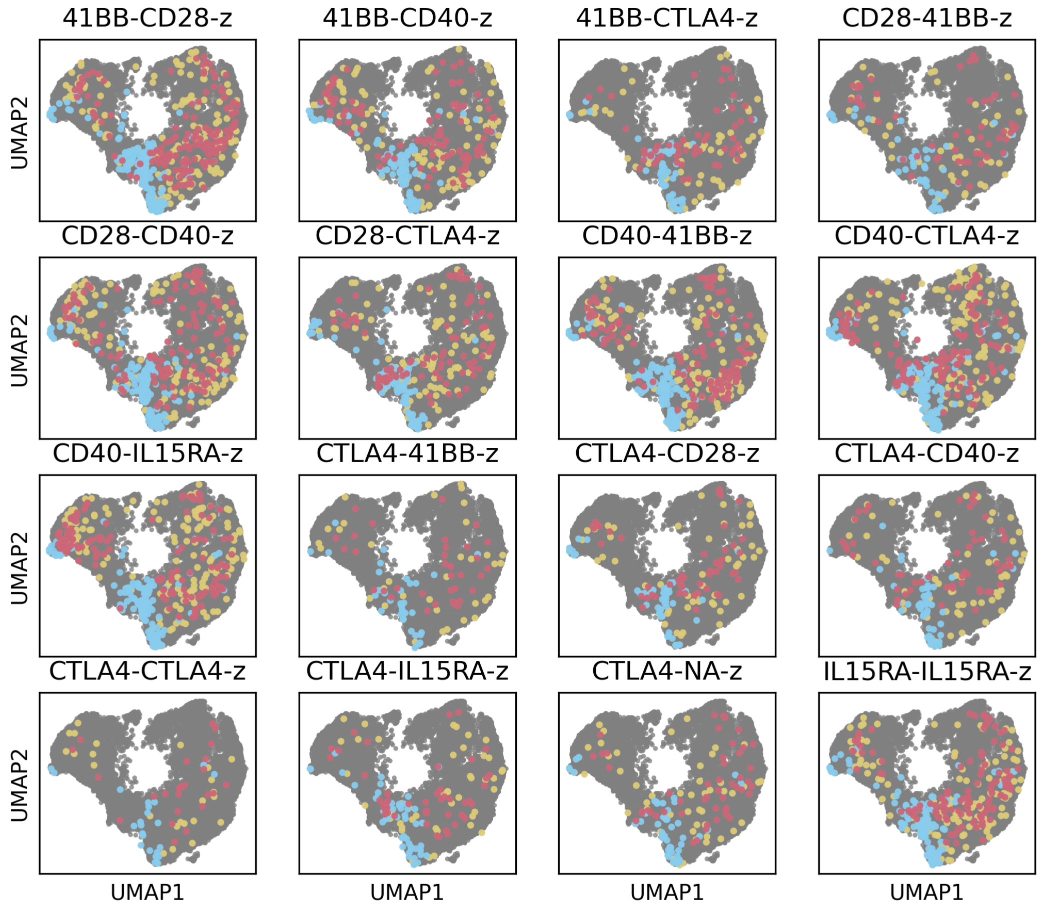


**Figure A9:** UMAPs of CD8 subset cells for the conditional OT model trained on CAR variants with >750 cells in the subset. All variants shown here were not present on in the training set, shown are all available cells for the variant

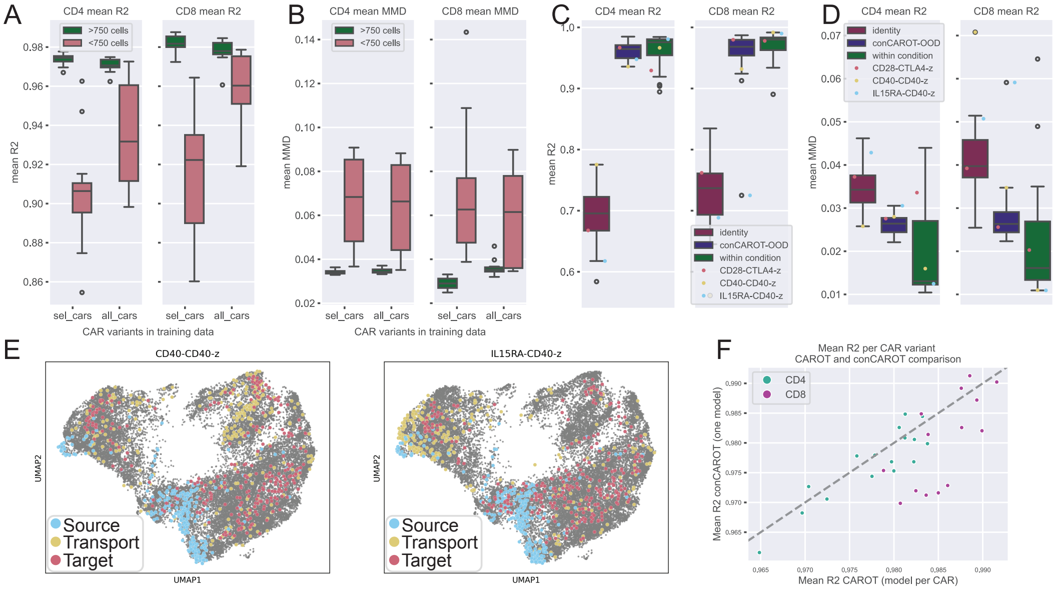


**Figure A10:** Additional OOD experiment results. A-B) Comparing performance of the model trained with all CAR variants (all_cars) and the model trained with only variants with >750 cells (sel_cars) (x-axis). The evaluation is split by CAR variants with >750 cells and variants with <750 cells. The variants with >750 cells are in distribution for both models, whereas the variants with <750 cells are OOD for the sel_cars model. Plots show the average score over nine samples per variant with the $R^2$ in A and the MMD in B. C-D) OOD performance of a model trained on all in-distribution CARs, leaving out one CAR at a time. Highlighted are CARs also shown in the UMAPs in the main text and in panel E that show a distinct response. Performance is again averaged over nine samples, with $R^2$ in C and MMD in D. E) UMAPs of CAR variants also shown in the main text for the OOD model evaluated in C&D. F) Comparison of CAROT (one model per CAR variant) and conditional CAROT (one model trained on all >750 cells CAR Variants), for all in-distribution CARs used in the conditional CAROT training. Variants for which conditional CAROT outperforms CAROT: CD4 - CD28-IL15RA-z, 41BB-CD28-z, IL15RA-41BB-z, CD28-CD40-z, and 41BB-CD40-z. CD8 - 41BB-41BB-z, IL15RA-41BB-z, and IL15RA-NA-z, CD28-NA-z

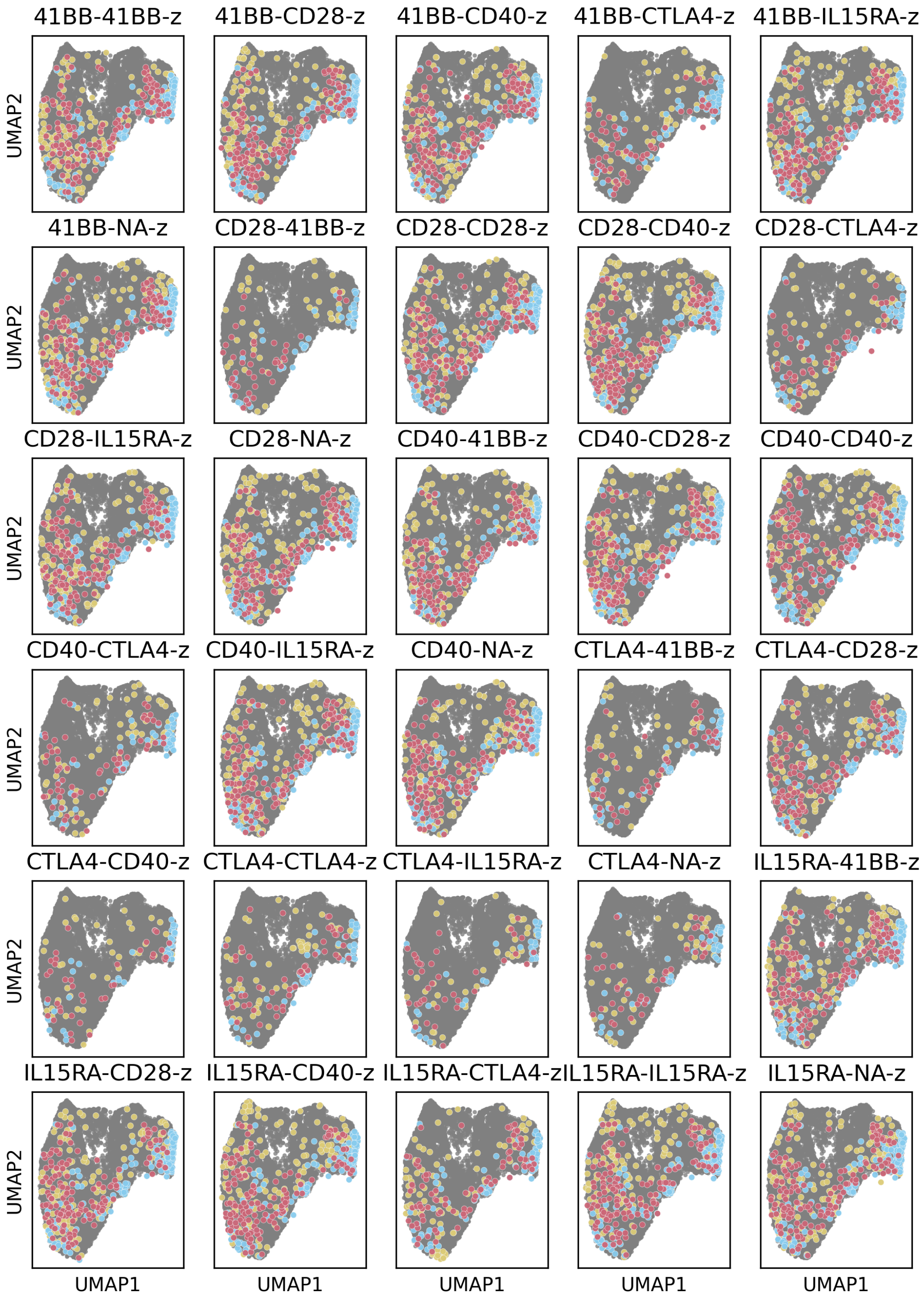


**Figure A11:** UMAPs of CD4 subset cells for the conditional OT model trained all CAR variants in the OOD setting, leaving the indicated CAR variant out during training.

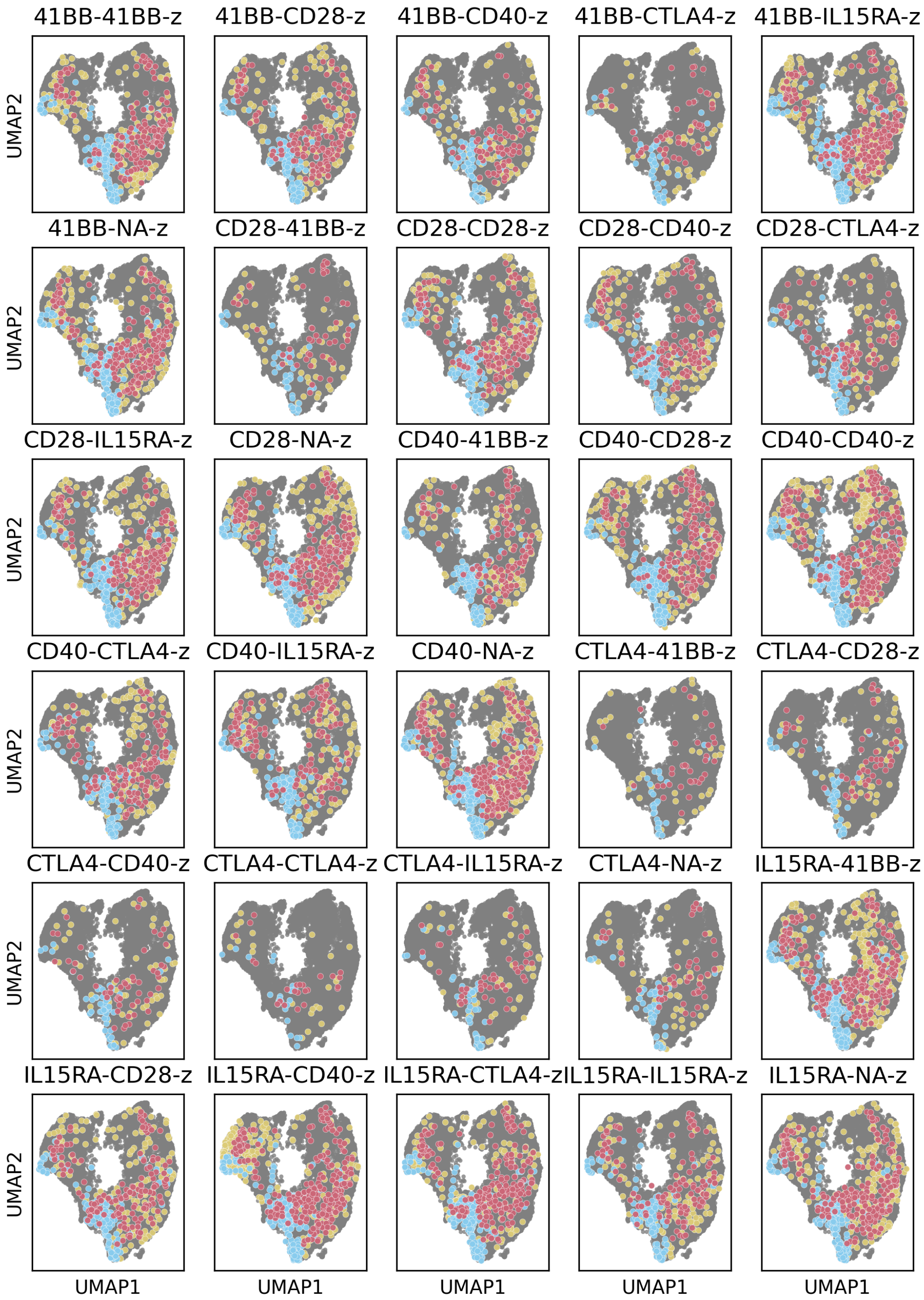


**Figure A12:** UMAPs of CD8 subset cells for the conditional OT model trained all CAR variants in the OOD setting, leaving the indicated CAR variant out during training.
